# Vectorchip: Microfluidic platform for highly parallel bite by bite profiling of mosquito-borne pathogen transmission

**DOI:** 10.1101/2020.10.19.345603

**Authors:** Shailabh Kumar, Felix J. H. Hol, Sujit Pujhari, Clayton Ellington, Haripriya Vaidehi Narayanan, Hongquan Li, Jason L. Rasgon, Manu Prakash

## Abstract

Mosquito bites transmit a number of human pathogens resulting in potentially fatal diseases including malaria, dengue, chikungunya, West Nile encephalitis, and Zika. Although female mosquitoes transmit pathogens via salivary droplets deposited during blood feeding on a host, very little is known about the genomic content of these nanoliter scale droplets, including the transmission dynamics of live pathogens. Here we introduce *Vectorchip*, a low-cost, scalable microfluidic platform for molecular interrogation of individual mosquito bites in a high-throughput fashion. An ultra-thin PDMS membrane coupled to a microfluidic chip acts as a biting interface, through which freely-behaving mosquitoes deposit saliva droplets by biting into isolated arrayed micro-wells enabling molecular interrogation of individual bites. By modulating membrane thickness, the device enables on-chip comparison of biting capacity and provides a mechanical filter allowing selection of a specific mosquito species. Utilizing *Vectorchip*, we show on-chip simultaneous detection of mosquito DNA as well as viral RNA from Zika infected *Aedes aegypti* mosquitoes – demonstrating multiplexed high-throughput screening of vectors and pathogens. Focus-forming assays performed on-chip quantify number of infectious viral particles transmitted during mosquito bites, enabling assessment of active virus transmission. The platform presents a promising approach for single-bite-resolution laboratory and field characterization of vector pathogen communities, to reveal the intricate dynamics of pathogen transmission, and could serve as powerful early warning artificial “sentinel” for mosquito-borne diseases.

## Introduction

Mosquito-borne pathogens and diseases including malaria, dengue, chikungunya, West Nile encephalitis, Japanese encephalitis, and Zika afflict more than 300 million individuals every year, resulting in more than 500,000 deaths [1]. As a result of increasing environmental pressures and human migration, these diseases are expected to continuously increase their geographical range, placing more than 50% of the world human population at risk within decades [2, 3]. To curb this crisis, we urgently need new tools to expand our understanding of mosquito-pathogen communities and transmission mechanics. In addition, expanding the scale of surveillance of infectious mosquito species and associated pathogens is critical for deployment of better-informed preventative measures in communities.

Molecular analysis of genomic material, proteins, or metabolites remains the gold standard for the detection and study of mosquito-pathogen communities in various settings. In order to obtain molecular samples for field ecology, mosquitoes are typically collected through different modalities of traps [4,5] or via human landing catches, which remain an ethically questionable method for gathering data [6–8]. These resource-intensive sample collection strategies commonly suffer from a limited scale of operation. As a result, severe undersampling of mosquito and pathogen populations remains a major concern around the globe [9]. In addition, molecular analysis of mosquitoes in the lab or field is usually performed using either (i) whole body sampling of mosquitoes by homogenizing them [10], (ii) analysis of mosquito saliva after extraction of salivary glands [11], or (iii) forced mosquito salivation [12] to determine the prevalence of pathogens in mosquito populations. In addition to requiring sample preparation of mosquitoes, these methods do not represent actual biting events thus providing limited information about the transmission dynamics such as infectious viral load in bites. More recently, alternative modes of sample collection exploiting e.g. sugar feeding or excreta have been proposed [13–17], but similarly do not simulate biting events, cannot differentiate between pathogens in saliva vs gut, provide population level (i.e. pooled) information with limited scope of improvement in resolution, and cannot quantify live viruses. A significant gap exists in our knowledge which requires novel tools to (i) quantitatively track the dynamics of pathogen transmission directly from single mosquito bites, and (ii) improve the throughput of molecular profiling of mosquito populations to boost surveillance strategies.

High-throughput molecular interrogation of saliva from mosquito bites can provide a new window into the dynamics of vector-borne disease transmission. Quantification of genomic content of salivary droplets and pathogen transmission directly from bites can help shed light on the dynamics of transmission at bite-by-bite resolution [18]. At the same time, parallel profiling of multiple biting events can lead to highly accelerated sampling of large vector populations in the field. Currently, an important barrier to developing such methods is the lack of a suitable vector-device interface. Membrane materials such as parafilm and animal casings typically used to interface with mosquitoes [19–21], lack the thermal and chemical compatibility needed to perform nucleic acid amplification, cell culture, and integration with microfluidic device manufacturing practices. As a result, it is imperative to look for novel materials and protocols that can change the landscape of vector-pathogen diagnostics.

In this work, we demonstrate *Vectorchip* - a microfluidic platform for molecular diagnostics of individual salivary droplets from mosquitoes allowing on-chip detection of mosquito DNA, viral RNA, and infectious viral particles directly from mosquito bites. This is enabled by a new fabrication strategy, resulting in reaction-ready, skin-mimic polydimethylsiloxane (PDMS) membranes which allow mosquitoes to bite and feed through them. We investigate the biting of various mosquito species using this chip, discovering biomechanical differences in biting capacity which can be used to create species-selective feeding barriers. Saliva from the biting activity of mosquitoes on *Vectorchips* is autonomously collected in individually isolated reaction chambers. Nucleic acid amplification assays conducted on-chip indicate that mosquito DNA and viral RNA are released during these biting events enabling identification of both the vector and pathogen. Finally, we show direct quantification of live, infectious viral particles from mosquito bites using focus forming assays (FFA) on chip, enabling interrogation of pathogen transmission through direct profiling of individual mosquito bites at high throughput.

## Results

### Device design and fabrication

Vectorchips with integrated PDMS elastomer membranes were fabricated using a combination of laser patterning and soft lithography. We selected PDMS as the material of choice due to its low cost, biocompatiblity, optical transparency, and chemical stability. PDMS-based devices can be fabricated using reproducible and scalable fabrication techniques [22]. Such devices have routinely been used for molecular analysis of small sample volumes using microfluidics [23], and are compatible with molecular assays and cell cultures [24, 25]. We used laser patterning to create open-ended arrayed compartments in PDMS for saliva capture (Fig. 1a-d). A thin elastomeric film was formed by spin-coating uncured PDMS over a flat silicon surface. We obtained membrane-integrated devices after plasma-induced PDMS-PDMS bonding of the laser-patterned array with the elastomeric film (Fig. 1e). Detailed steps describing chip development are included in the methods section. The fabrication process is extremely versatile and allows for user-defined variation in the size and density of the individual compartments. We were able to fabricate chips with hole diameters ranging from 150 μm to 1 cm (Fig. S1) showing the scales of operation for the laser-ablation process while maintaining membrane stability. Additionally, the membrane can be easily integrated with fluidic networks for direct interfacing with mosquito bites, enabling assays involving on-chip fluid exchange (Fig. 1f). The microfluidic compartments on the chips can hold feeding media such as blood or sugar water (Fig. 1g), collect saliva during biting events, and act as isolated reaction chambers for molecular assays.

**Figure 1:**
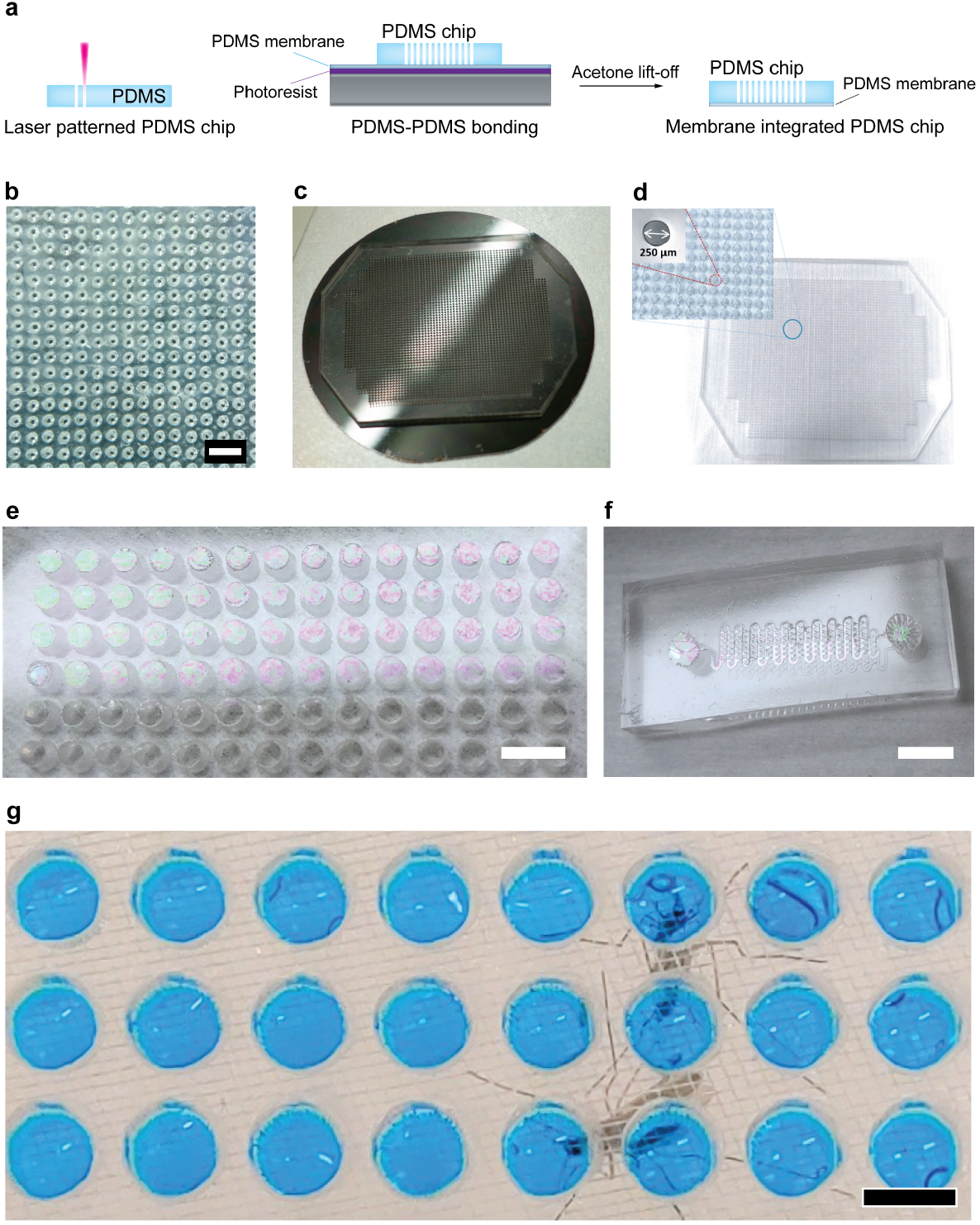
Fabrication of *Vectorchips* for collection of mosquito saliva and molecular assays. a) Schematic showing the fabrication process of *Vectorchips*. Laser patterning is used to obtain through-holes in blocks of PDMS. A thin sacrificial layer of photoresist is spread onto a silicon wafer followed by spin-coating a thin film of PDMS on the wafer surface. The laser-patterned body of the chip is then bonded to the thin film of PDMS using plasma activated PDMS-PDMS bonding. Membrane-bonded chips are obtained by placing the wafer in an acetone bath, where the sacrificial layer of resist is dissolved. (b) A PDMS block with laser-generated through-holes (diameter 250 μms). (c) A PDMS chip bonded to a 1.6 μm thin PDMS membrane on a 4 inch silcon wafer. (d) A *Vectorchip* with around 3000 wells (dia: 250 μms) is shown. (e) Chip with well diameter 1.75 mm and a visible PDMS membrane (thickness 1.6 μm). (f) Design fluidity in fabrication allows production of channel-integrated chips with a PDMS membrane on top. (g) These wells can be loaded with feeding solution, reaction reagents, and can store mosquito saliva after biting assays. Wells loaded with feeding media (10% sucrose laced with blue food color) shown here. Scale bars represent 1 mm in (b), 5 mm in (e), and 2 mm in (f, g).

### Mosquito-chip interactions

We tested the ability of mosquitoes to pierce through the PDMS membranes. Mosquitoes were attracted to the chips using heat as a guiding cue. We placed a camera above the chip and observed stylet insertion through 1.6 μm thick PDMS membranes (Fig. 2a, Supp. Movie 1,2, Note: thickness of the PDMS membrane at 1.6 μm was selected by maximum spin speed available at the time of these experiments). Abdominal engorgement in mosquitoes was observed after biting, indicating that mosquitoes can successfully feed through these membranes. Four species of mosquitoes were tested (*Aedes aegypti, Aedes albopictus, Culex tarsalis, Culex quinquefasciatus*) and demonstrated successful probing and feeding through 1.6 μm thick PDMS membranes.

**Figure 2:**
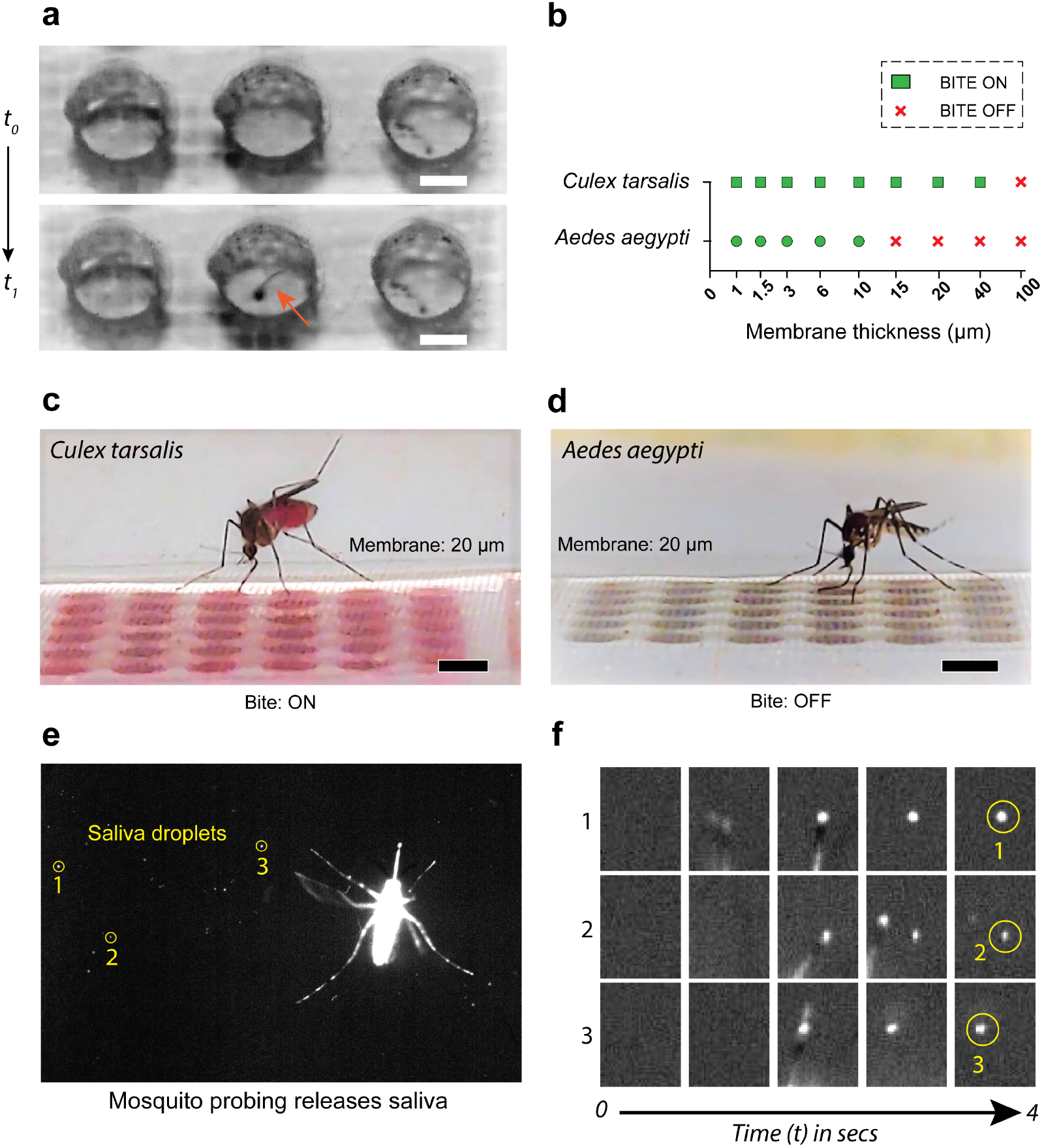
Mosquito biting on *Vectorchips*. (a) Stylet entry through a PDMS membrane for an *Aedes aegypti* female. (b) Mosquito feeding success as a function of membrane thickness for two mosquito species - *Aedes aegypti* and *Culex tarsalis*. We demonstrate that by tuning the membrane thickness we can turn biting “on” or “off”. Biting “off” is defined as when no mosquito in the cage was able to feed from the chip in a duration of 30 minutes. (c) A *Culex tarsalis* female mosquito biting through a 20 μm thick PDMS membrane. (d) Bending of the proboscis observed for an *Aedes aegypti* female, indicating failure in biting through the 20 μm membrane. (e) Fluorescent salivary droplets expectorated by an *Aedes aegypti* mosquito while probing. These mosquitoes were fed on rhodamine-laced sugar water resulting in fluorescent saliva. (f) Timelapse images show a magnified view of fluorescent salivary droplet deposition over a period of 4 seconds. Scale bars represent 500 μms in (a) and 2 mm in (c, d).

We also examined whether the membrane thicknesses can be engineered to selectively prevent biting of some species while letting others feed. We loaded the laser-patterned microfludic compartments with blood and warmed the chips to attract mosquitoes (Fig. S2, Supp. Movie 3). We considered biting to be “off” when none of mosquitoes in the cage demonstrated abdominal engorgement within 30 minutes since start of the experiment. Tests were performed with *Aedes aegypti* (~ 45 mosquitoes per cage) and *Culex tarsalis* (~ 25 mosquitoes per cage) while varying the thickness of the PDMS membrane from 900 nm to 100 μms. We observed significant differences in the biting ability of the two species, where complete inhibition of feeding in *Aedes aegypti* was observed with membrane thickness at or greater than 15 μms (Supp. Movie 4, 5). In contrast, complete inhibition of feeding with *Culex tarsalis* was observed for membranes that were 100 μms thick (Fig. 2b). Even though a larger population of *Aedes aegypti* were available to feed from the chips, their biting strength appeared to be inferior to *Culex tarsalis*. This remarkable difference in probing capacity of these two species has not been reported earlier. While some studies have focused on understanding the biting mechanics of a single mosquito species (*Aedes aegypti* or *Aedes albopictus*) [26, 27], we are not aware of any study which looks at the difference in biting strength between different species and the evolutionary or biomechanical differences involved in this variation. This intriguing biomechanical phenomenon can be advantageous by deploying devices where membranes of desired thickness can act as species-selective filters, providing a means to better track highly relevant species and pathogens in the field.

In order to extract molecular information from mosquito bites, the chip aims to exploit the release of saliva during probing and biting events. The release of salivary droplets can be visualized using fluorescence imaging of mosquitoes fed on rhodamine-labeled sugar water, which renders the saliva fluorescent (Fig. 2e, Supp. Movie 6). Time sequences (Fig. 2f) show magnified images of three locations on a membrane surface where short probing activity results in deposition of salivary droplets. These salivary droplets house several biomarkers in the form of mosquito salivary proteins [28], active pathogens, mosquito cells and/or nucleic acids that can be used for their identification [12, 18, 29].

### Tracking dynamics on-chip

A major consideration for biting based diagnostics would be the ability to sample either the whole population or individual mosquitoes. The relationship between number of mosquitoes interacting with each isolated reaction chamber should depend on the number of mosquitoes in the cage, the time of interaction, number of wells, and the area of the chip. It is possible that the feeding media also affects the number of biting events performed by a female mosquito. We utilize sucrose as the feeding media since it is a diet source that does not inhibit polymerase chain reaction (PCR) assays and therefore will limit the analysis to this feeding solution. In order to perform sucrose biting assays for molecular diagnostics, we filled the wells with 10% sucrose, and covered the open side of the chip with a glass slide. A resistor which acts as a heat source was placed atop the glass slide and the chip-resistor assembly was placed on top of the mesh ceiling of a mosquito cage. A part of the chip was protected by a paper tape such that mosquitoes cannot bite into the membrane. The area of the chip protected by tape serves as negative control for the biting assays. A camera was placed on the bottom of the cage to track and record mosquito activity on chip (Fig. S3). Image recognition algorithms were used to analyze the obtained images and track the presence and movement of mosquitoes on the chip (Fig. 3a-d, Supp. Movie 7). Tracking data reveals the dynamics of mosquito-chip interactions indicating the trajectories followed by mosquitoes and the time spent by mosquitoes at different spots on the chip. Regions with non-overlapping trajectories reveal the areas which are most-likely to be probed by a single mosquito (Fig. 3d).

**Figure 3:**
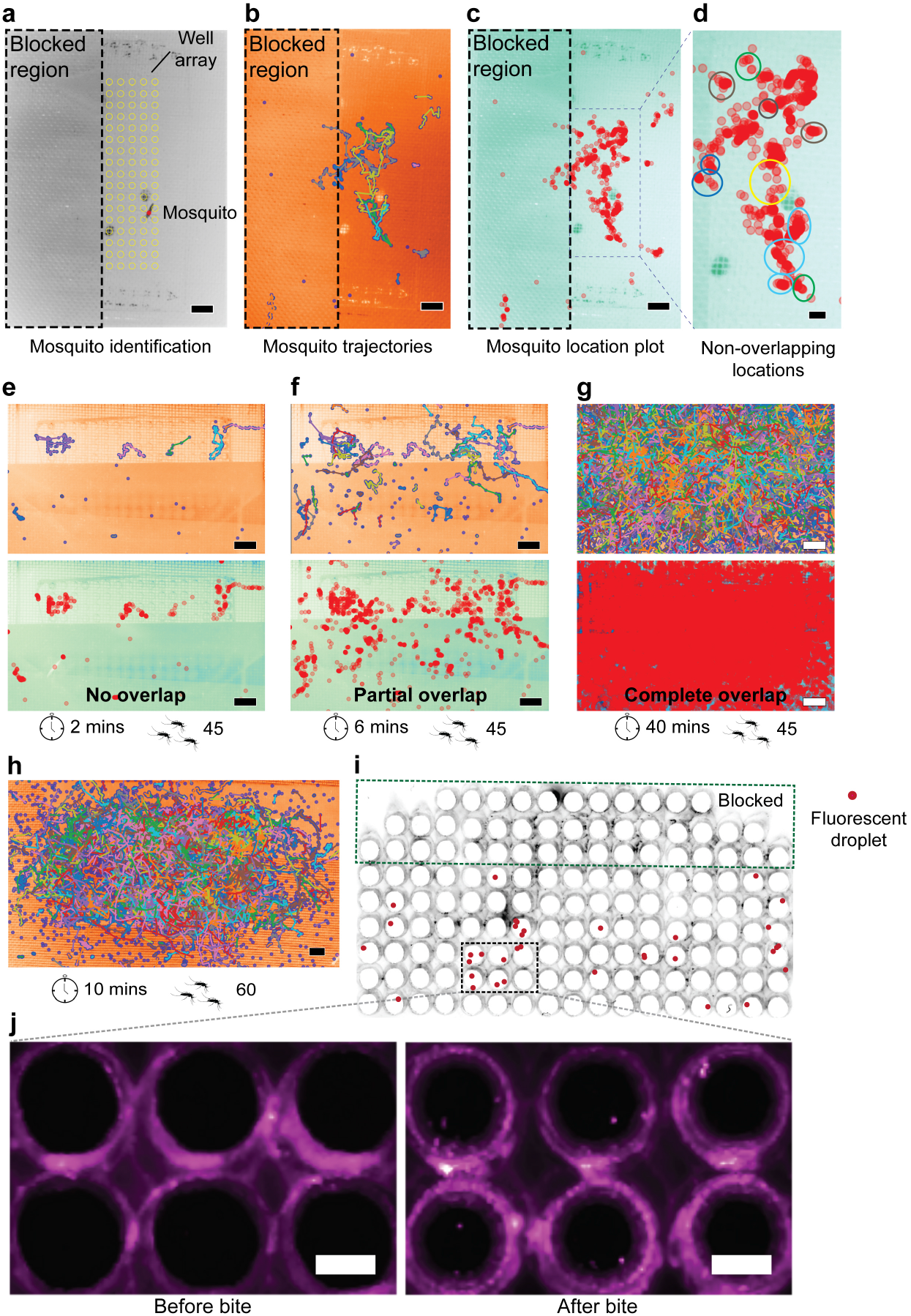
Tracking mosquito activity on chip. (a) Image recognition software is used to identify and track mosquitoes on the chip. Wells are highlighted in yellow. (b) Trajectories of mosquito movement can be plotted and indicate the overlap between different movement tracks. (c) Location plot of mosquitoes on the chip indicated by red circles. Regions with overlapping red circles are darker in color. (d) Trajectory and location plot can be used in combination to identify regions with individual mosquito locomotion activity as compared to regions with more crowding. (e-g) Varying experimental parameters (time of biting shown here) can be used to obtain either individual mosquito plots or population data. Three cases are discussed where there is (e) no overlap between multiple probing trajectories, (f) partial overlap and (g) complete overlap. (h-j) Tracking and corresponding salivary droplet deposition data for a chip. (h) shows extensive translocation activity of mosquitoes on chip. (i, j) Several wells show single fluorescent droplet indicating probing by a single mosquito, whereas other wells show multiple probe marks. The tracking and probing data suggest that we should be able to obtain both individual bite data as well as multi-bite statistics on *Vectorchip*. Scale bars represent 5 mm in (a-c, e-h), 2 mm in (d), and 1 mm in (j).

On-chip location and tracking dynamics for mosquitoes on different chips is shown in Fig. 3e-g and Fig. S3. Three cases are demonstrated - i) over a short time period and low mosquito density, no overlap between any of the trajectories is found (Fig. 3e). ii) Increasing the time of interaction, we can simultaneously obtain both individual tracks and pooled probe data (Fig. 3f). iii) An extended interaction time between mosquito population and the chip results in complete overlap between trajectories (Fig. 3g). The results indicate that by modulating the time of interaction between the mosquito population and the chip, the user can control the degree of probing experienced by the chip. As an example, for a chip where there is no overlap between trajectories, every probed well should provide single mosquito-resolution data.

While the tracking of mosquitoes on chip provides information about mosquito locomotion and time spent over different areas of the chips, fluorescent salivary emission can be used as a means to gauge the extent of probing over different regions of the chip. We quantified the number of spots seen in different wells of the PDMS chip by using fluorescent scans of the chip before and after biting (Fig. 3h, i). No spots were observed in the negative control region where access was blocked to mosquitoes. Several spots were observed on the open areas of the chip, with several wells indicating that they have been bitten single time. These datasets indicate that the *Vectorchip* provides opportunities to obtain data at both the individual bite level as well as pooled-profiling of multiple bites.

### PCR diagnostics on-chip

We selected devices with PDMS membrane thickness of 1.6 μm and well diameter of 1.75 mm as the first prototype to perform molecular analysis of salivary samples. These well dimensions were amenable with manual pipetting while providing a dense array of 150 independent reactions per chip (within a chip area of 5 cm × 2.5 cm).

Detection of mosquito mitochondrial DNA (mtDNA) and RNA from viruses in saliva was performed using on-chip PCR with end-point fluorescence readout. A detailed protocol for performing PCR in *Vectorchips* is provided in the Methods section and Supporting Information file (Fig. S4). We tested the efficiency of PCR amplification in *Vectorchips* by manually loading a known concentration of DNA and RNA into the reaction wells (Fig. 4a-c). We performed the PCR for 42 cycles and detect DNA amplification from approximately 5 DNA copies in 4 μL of reaction mix (~ 1 copy/μL) (Fig. 4b). Simultaneously for Zika reverse transcription-PCR (RT-PCR), viral RNA detection was successfully demonstrated from approximately 150 RNA copies in 4 μL of reaction mix (Fig. 4c). We verified the performance of end-point PCR in vectorchips by running reactions with identical reagents, volumes and conditions in 96 well plates. Both qPCR amplification curves and end point fluorescence information was obtained from the well plates. The reactions were tested in DI water and also in the presence of dried sucrose (10%) to ensure that amplification response correlated well with manually spiked concentrations and the presence of sucrose did not inhibit the reactions (Fig. S5(a,b)).

**Figure 4:**
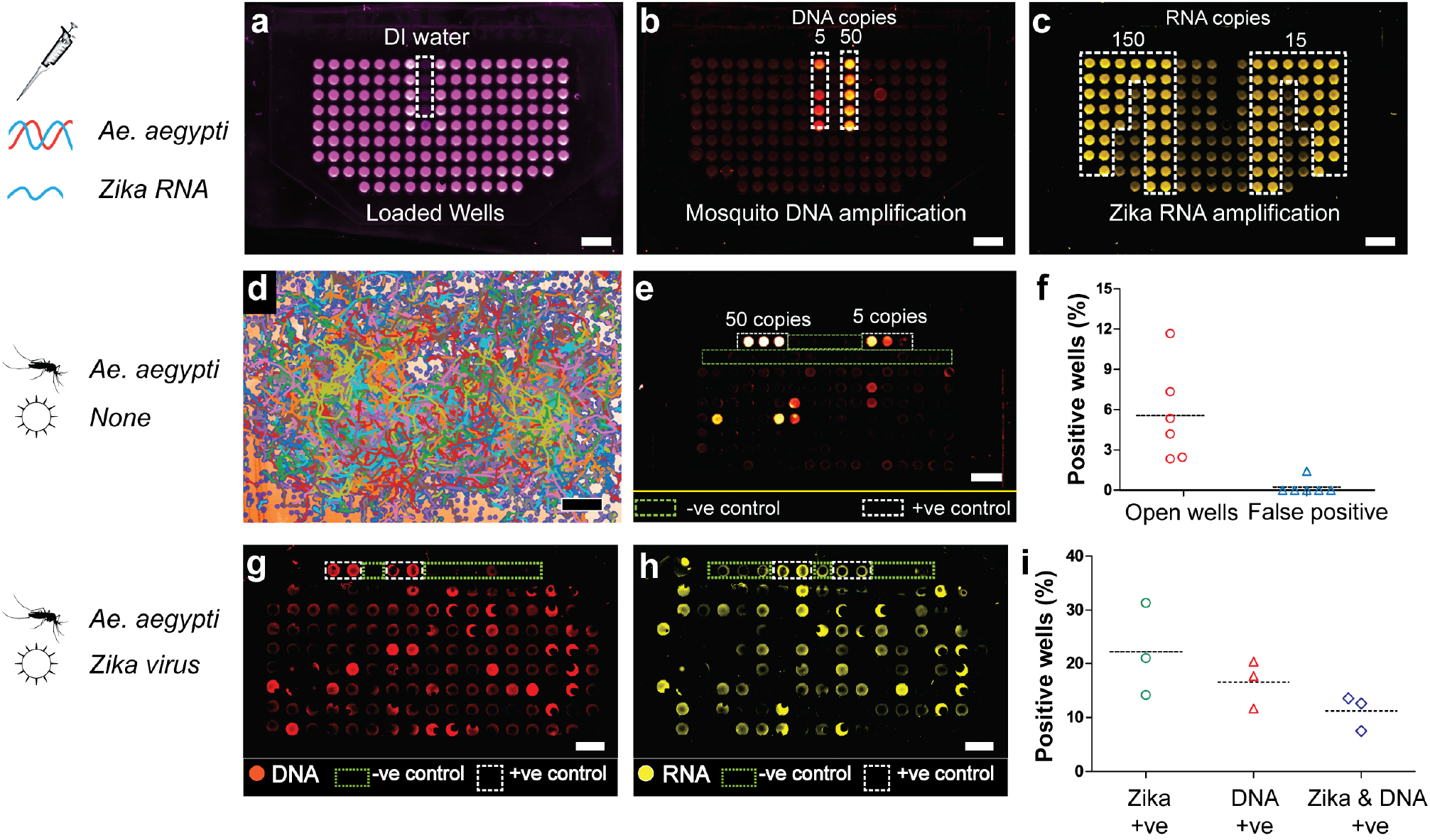
On-chip PCR for detection of mosquito and pathogen. (a-c) A spiked assay demonstrates successful amplification of *Aedes aegypti* DNA and Zika RNA on chip. (a) indicates wells filled with PCR mix, with ROX dye providing the background fluorescence. Fluorescent wells indicate amplification of (b) mosquito DNA, and (c) Zika RNA on chip. Assays were performed with uninfected mosquitoes where (d) tracking patterns were collected and demonstrate extensive translocation activity on the chip. (e) PCR shows detection of mosquito DNA on the chip directly from biting. (f) Percentage of wells available for biting on *Vectorchip* that display a positive PCR outcome. The false positive rate (amplification in wells that were covered by tape) was close to zero (0.23%). Assays performed with Zika infected mosquitoes indicate presence of both (g) *Aedes aegypti* DNA and (h) Zika virus RNA after bites on chip. (i) Percentage of wells available for biting that show a positive signature for Zika RNA, mosquito DNA or both. Scale bars represent 5 mm in (a-e) and 2 mm in (g, h).

The chips were allowed to be probed by mosquitoes for 20-45 minutes. While testing uninfected mosquitoes, PCR assays were performed on-chip for detecting the presence of mosquito DNA in deposited salivary droplets. Fig. 4d-e shows the tracking data and PCR detection of mtDNA for a chip which was placed on a cage with 75 uninfected *Aedes aegypti* females for 45 minutes. Areas highlighted in green were protected by a paper tape such that mosquitoes cannot probe them and serve as negative controls. The tracking and PCR data obtained from the chip indicate significant locomotive activity of the mosquitoes on-chip, and detection of mosquito DNA in wells. Multiple experiments indicate that a subset (3 - 12%, n = 6) of the wells where mosquitoes are active test positive for presence of mosquito DNA (Fig. 4f). The rate of false positives obtained from wells not accessible to mosquitoes was very close to zero (0.23%, n = 6). In order to further validate the rate of detection of mosquito DNA on *Vectorchips*, we performed a biting assay using a 96 well plate loaded with agarose and covered with a parafilm membrane. We recorded the tracking and DNA amplification response from uninfected mosquito bites on the well plate (Fig. S5(c,d)). Around 15% of the wells where mosquitoes were active tested positive for presence of mosquito DNA, which is close to the fractions observed on vectorchips. Since mosquito DNA can only be detected in a subset of wells showing mosquito activity, this suggests that the presence of detectable levels of mosquito DNA in saliva is a noisy physiological process, likely dependent on multiple variables such as duration of feeding and time of last bite. It is furthermore important to note that our tracking algorithm only detects the presence of mosquitoes, but does not provide information regarding if a well was bitten or not.

Fig. 4h shows results obtained when *Aedes aegypti* infected with Zika virus [30] interacted with the chip for 20 minutes. We subsequently performed RT-PCR assay on the chip demonstrating that *Aedes aegypti* mtDNA and ZIKV RNA can be detected in 24/118 and 37/118 wells, respectively. We summarize the detection rates of mosquito DNA and Zika RNA for three assays performed with ZIKV-infected mosquitoes in Fig. 4i. Interestingly, not all wells positive for ZIKV RNA show positive for mosquito DNA and vice versa, highlighting the stochastic composition of bite-derived saliva droplets. A higher fraction of wells (mean ~ 22%, n = 3) showed positive for ZIKV RNA as compared to mosquito mtDNA (mean ~ 17%, n = 3), indicating a higher prevalence of viral genomic material in vector saliva as compared to mosquito mtDNA. Detection of mosquito DNA and virus RNA in these assays demonstrates the capacity of this tool towards the identification of mosquito and pathogen species directly from mosquito bites on-chip.

### Detection of infectious viral particles directly from bites

While PCR-based end-point assays are an important strategy for multiplexed detection of vectors and pathogens in various settings, they cannot determine the presence or concentration of infectious virus particles in bites. Detection of infectious pathogen load in mosquito bites is important to understand the vector competence of mosquito-pathogen systems and its dependency on factors such as local climate, mosquito physiology [31], and presence of endosymbionts (e.g. *Wolbachia*) [32, 33]. Typical methods to measure vector competence rely on manual “forced salivation” of individual mosquitoes [10], where the viral loads may differ from biting events. We tested the *Vectorchip* to perform focus forming assays (FFA) and detect infectious viral particles directly as a result of mosquito bites on chip (Fig. 5).

**Figure 5:**
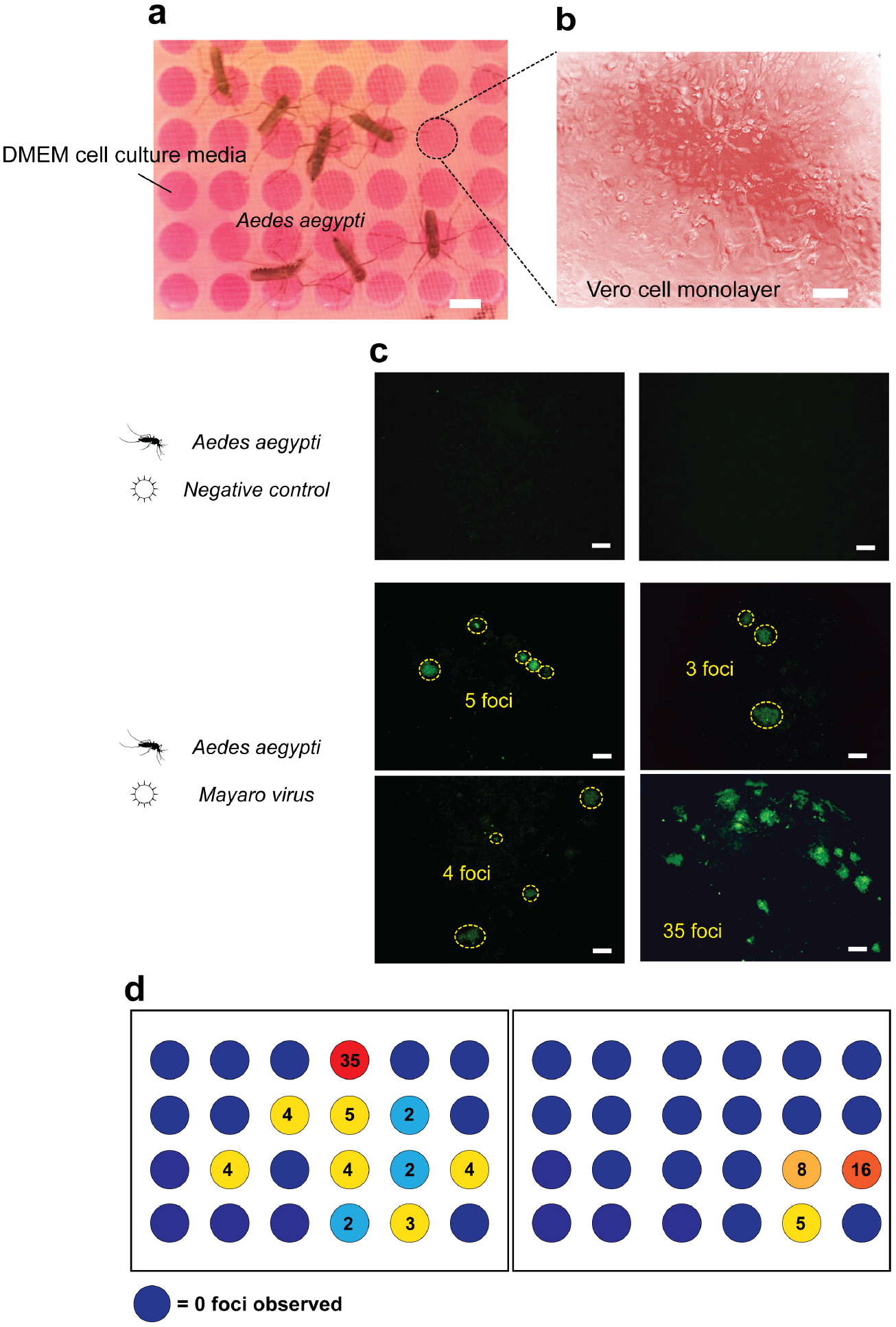
Quantifying active viral particles using focus forming assays. Focus forming assays on *Vectorchips* can directly quantify the number of active viral particles in mosquito bites, without the need for isolation of individual mosquitoes and manual salivary extraction. (a) Image shows *Aedes aegypti* mosquitoes infected with Mayaro virus biting on *Vectorchips* filled with DMEM cell culture media. (b) A monolayer of vero cells growing on *Vectorchip* membrane. (c) FFA performed on a *Vectorchip*. Control samples and compartments with active Mayaro virus are shown. Green channel shows fluorescence from an antibody against a viral envelope protein. Every green island represents an active viral particle. No active viral particles were seen in the control wells. (d) FFA formed on two chips and viral foci were counted using the antibody fluorescence. Scale bars represent 2 mm in (a), 30 μms in (b) and 100 μms in (c).

The wells were filled with cell culture media (Dulbecco’s modified eagle medium - DMEM). The efficiency of FFA on-chip was tested initially using manual spiking of viral particles into *Vectorchips* (Fig. S6). In order to perform biting assays, *Aedes aegypti* mosquitoes were infected with Mayaro virus, an emerging zoonotic pathogen which causes a dengue like disease [34]. The mosquitoes were able to bite through the membrane into the wells and transfer live Mayaro viral particles directly into the cell culture medium (Fig. 5a). These viral particles then infect a monolayer of vero cells cultured on the PDMS membrane. These infections can be identified via fluorescent antibodies specific to viral envelope proteins (Fig. 5c). Each fluorescent patch (focus) is attributed to infections transmitted by a single viral particle. Foci counted on two sample chips can be seen in Fig. 5d. No foci were visible from negative control samples. The distribution of viral foci on the chips indicate the heterogeneity of infectious particle dose in mosquito bites. The data in Fig. 5 demonstrate the suitability of using the *Vectorchip* to probe the transmission of infectious viral particles.

## Discussion

We demonstrate a novel microfluidic device integrated with a PDMS “ artificial skin” membrane for molecular diagnostics of individual mosquito bites on a chip. The device permits user-desired modifications in chip design as well as more than two orders of magnitude variation in reaction chamber volume for performing molecular analysis. The elastomer membrane is compatible with scalable device fabrication, microfluidic integration, nucleic acid amplification assays, as well as cell culture assays, allowing multimodal interrogation of mosquito bites on chip. Interfacing mosquito bites with fluidic channels provide the potential of using this device with integrated vascular pathways or particle traps to study transport and subsequent infection dynamics of expectorated pathogens in a user-controlled environment. As compared to previous methods, the *Vectorchip* uniquely allows quantitative interrogation of mosquito bites on a chip, which may answer questions such as the genomic diversity of bites, transmission dynamics of pathogens in bites, and enable high throughput diagnostics of large mosquito populations at single bite resolution.

We tested our devices with four mosquito species and all were able to probe through ultrathin PDMS membranes (thickness 1.6 μms) demonstrating the ability of this device to interface with variety of vectors. Furthermore, our novel fabrication strategy permits us to engineer the “skin characterstics” such as thickness and stiffness. We discover that different species of mosquito can display significant variation in their biting strength, the biomechanical basis for which remains an intriguing question. This variation in probing capacity provides an opportunity to perform mechanical species selection for on-chip sample collection by careful engineering of the membrane thickness and stiffness. This observation also provides opportunities towards examining the biting capacity in different mosquito species and if there exists an evolutionary relationship of the biting ability with their preferred prey species.

We also utilize on-chip tracking of mosquito activity to demonstrate that varying the experimental parameters including chip design and number of mosquitoes can allow collection of bite data from the population as a pool or from individual bites. While population data can broadly indicate the presence of infectious agents in the community, individual bite statistics can help explore these phenomena at a higher (bite-by-bite) resolution.

The *Vectorchip* aims to significantly reduce the manual labor, time, and risk associated with capture, individual segregation and sample extraction from individual mosquitoes as traditionally practiced. Reaction in miniaturized chambers reduces the cost of molecular assays - chips used in this report (1.75 mm diameter) performed PCR reactions at 30 ¢per reaction chamber which is approximately 5× lower than current single mosquito homogenization assays. We also fabricated further miniaturized device designs with smaller well diameters, which can provide a 100× reduction in individual reaction volumes and lower the cost per-assay by two orders of magnitude (*<*1 ¢per reaction).

We detect that biting mosquitoes release their DNA on chip, which can be used to identify their species as well as targeted genetic mutations. Detection of these genetic signatures can be used for monitoring the spread of native or invasive mosquito species globally, insecticide resistance in mosquitoes, or tracking the range of gene-drive mosquitoes released at experimental sites [35]. Detection of pathogens was demonstrated using i) RT-PCR assays for the presence of viral nucleic acids and ii) FFA to quantify infectious viral particles directly from mosquito bites. PCR assays can enable low-cost sampling of pathogens in the field informing community action for disease prevention. Simultaneously, FFA results provide a window to dynamics of infection transmission directly through individual mosquito bites, which as of yet remains difficult to study.

Currently, ecological distribution of pathogens in our environments is significantly undersampled. This contributes to dramatic consequences in the form of unexpected viral outbreaks. There is an urgent need for vector-pathogen sampling to shift from current resource-intensive practices to low-cost technologies that can perform large-scale surveillance. We hope tools like the *Vectorchip* will allow us in the future to completely eliminate human-landing catches. The *Vectorchip* provides a route for multimodal analysis of vector-pathogen dynamics both in lab and field including vector identification, behavior and pathogen surveillance at a high throughput and resolution. We anticipate the use of *Vectorchip* tool can be expanded from vector and pathogen surveillance testing in field sites and lab settings to study of vector biting behavior on chip and expand to other biting vectors such as ticks and sandflies.

## Methods

### A) Chip fabrication

#### a) Main body

PDMS chips with thickness ~ 4 mm were prepared as the main body of the chip. Sylgard 184 base and cross linker (Dow Chemicals, USA) were mixed at a 10:1 ratio and cured at 75 °C on a flat heater surface. Laser ablation (60-Watt Mini CO_2_ laser, Epiloglaser, USA) was used to cut holes through the PDMS blocks with desired hole diameter and density. Debris produced by the laser ablation process was removed using 1 hour sonication in acetone. The chips were then rinsed with DI water and dried.

#### b) PDMS membranes

PDMS membranes were prepared over single side polished, 4 inch silicon wafers. A layer of positive photoresist (Microposit™ S1813G2) was spun on the wafers to a thickness of around 2 microns (1500 rpm for 45 s). The resist was softbaked at 80 °C for about 30 minutes. PDMS (Sylgard 184 kit with base and curing agent mixed at 10:1 ratio) was freshly prepared and spuncoat on top of the wafers. The thickness of the PDMS coat was controlled by varying the spin speed and time. In order to obtain a membrane with thickness close to 1.6 microns, PDMS was spun on the wafers at 6500 rpm for 10 minutes. Wafers with thin layers of PDMS were cured at 70 °C for at least 6 hours. The thickness of the PDMS on the wafers was measured by cutting strips off of a silicon wafer and measuring the difference in height using a profilometer (KLA Tencor Alpha Step D500).

#### c) Vectorchips

The main body of the chip and wafers with thin PDMS membranes were placed in a plasma cleaner and exposed to O_2_ plasma (80 W, 45 s, 2 sccm O_2_ flow). The PDMS chips were then placed in contact with the thin PDMS membranes to enable bonding of the membranes to the chips. The wafers were baked at 80 °C for around 2 hours to improve the bonding strength. A razor blade was used to cut outlines around the chips, such that membrane-attached PDMS chips can detach from the wafers. The wafers were then placed in acetone for approximately 2-4 hours. The sacrificial resist layer is dissolved in acetone and PDMS chips with integrated ultrathin membrane are separated from silicon wafers. The chips are then rinsed gently in DI water and dried in an oven.

### B) Biting assays

Vectorchips were hydrophilized by placing in a plasma cleaner (Harrick Plasma) at high power for 5 mins.

In order to test the biting ability of mosquitoes, sugar water was removed from the cages approximately 18 hours prior to the assays. Chips with varying membrane thickness were filled with defibrinated sheep blood (Hemostat laboratories, USA). The well diameter was kept constant at 1.75 mm. Chips were placed on mosquito cages for approximately 30 minutes, and blood feeding was monitored visually.

For PCR assays, the wells were filled with 10% sucrose solution. Sugar water was removed from the mosquito cages around 6-18 hours prior to the biting tests. The chips were then placed on top of a mosquito cage. A strip of tape was adhered on top of the cage and the chip was placed such that some rows were placed atop the tape and were thus inaccessible for the mosquitoes to bite through. A heat source (resistive heater or tissue culture flask with warm water) was placed atop the chips such that their temperature was approximately 37 °C. Mosquitoes bite the chips for approximately 20-45 minutes, expectorating saliva in the process which is collected in the wells. A raspberry pi camera attached to the bottom of the cage records mosquito activity on-chip. After approximately 20-45 minutes, the chips are removed from the cages. Chips bitten by ZIKV infected mosquitoes were placed on a hotplate set to 98 °C for 15 minutes to inactivate the virus and facilitate downstream processing.

PDMS base (Sylgard 184 part A) and crosslinker (Sylgard 184 part B) were mixed to a weight of 1 gram and placed in a vacuum degasser for removing any bubbles. A glass slide (size: 75 mm × 50 mm) was taken and approximately 0.75 g of the freshly prepared PDMS was added to the slide. The PDMS was spread evenly using a pipette tip and was degassed for 2 minutes to remove any bubbles. The *Vectorchip* used for biting assay was then placed on the slide, membrane-side down into the thin layer of uncured PDMS carefully.

The chip was then dried either at room temperature overnight or in an oven at 80 °C for 20 minutes (for Zika-infected mosquitoes). This process adheres the chip to a solid glass slide by curing of the thin PDMS layer and allows the sugar water to evaporate in every well.

### C) Mosquito tracking algorithms

Image recognition algorithms were written in Python 3 see ref. [21] for details, and used to track mosquito movement on the chip and are available on github (https://github.com/felixhol/vectorChip).

### D) RT-PCR assays

TaqMan™ Fast Virus 1-Step Master Mix was used for the amplification assays. Following primers and probes were used for the amplification of *Aedes aegypti* mitochondrial DNA and Zika RNA.

*Aedes aegypti* Fwd Primer: ACACATGCAAATCACCCATTTC
*Aedes aegypti* Rev Primer: CATTGGACAAGGCCTGTAACT
*Aedes aegypti* mtDNA probe: HEX-AGCCCTTGA-ZEN-CCTTTAACAGGAGCT-3IABkFQ
Zika virus Fwd Primer: CCGCTGCCCAACACAAG
Zika virus Rev Primer: CCACTAACGTTCTTTTGCAGACAT
Zika virus probe: FAM-AGCCTACCT-ZEN-TGACAAGCAGTCAGACACTCAA-3IABkFQ

The RT-PCR reaction mix was prepared as follows:

For every 20 μL of reaction mix:

Fast virus mix - 5 μL
Fwd Primer - 1 μL
Rev Primer - 1 μL
Fluorescent probe - 0.5 μL
DI water - add as required to total 20 μL

Four μL of reaction mix was pipetted into all wells. Positive control wells were filled with *Aedes aegypti* DNA or Zika virus RNA. Silicone oil (Sylgard 184 part A) was poured over the wells to prevent against evaporation and allowed to penetrate the wells. If needed, gentle degassing for 15 minutes helped the oil enter the wells. The chips were then placed on a flat-plate thermocycler (GenePro, Bioer Tech.) and RT-PCR was performed for 42 cycles.

50 °C: 10 mins
95 °C: 45s
cycle × 42 (95 °C: 35s; 60 °C: 80s)

The chips were then scanned using a fluorescence image scanner (Typhoon FLA 9000, GE Health-care) to obtain the images.

qPCR tests (StepOnePLus, Applied Biosystems) were performed in 96 well plates using identical samples and conditions as the *Vectorchips* in order to validate the RT-PCR efficiency in the chips.

### E) Focus forming assays

Chips were loaded with DMEM cell culture media in each well and placed over cages with approximately 50 female Mayaro virus infected *Aedes aegypti* mosquitoes. The mosquitoes were allowed to bite the chips for 30 minutes. The negative control chips were bitten by uninfected mosquitoes. Chips were then removed from the cages and 10 μL of Vero cell suspension (1000 cells/μL) was dispensed into each well. To allow the virus particles to adsorb on to the cell surface, chips were incubated for one hour at 37 °C incubator with 5 % CO_2_. To remove unbound viral particles, cells were washed once with DMEM without FBS. Thereafter, 20 μL of 1% methylcellulose (MC) overlay medium supplemented with DMEM and 5% FBS was added to each well of the chips. Chips were then incubated at 37 °C incubator with 5% CO_2_ for 24 h. Note, to minimize the evaporation rate of culture media from the wells the chips were incubated in a humidified secondary container. The MC overlay medium was removed, and cells were fixed with 4 % PFA for 20 minutes at room temperature (RT). After the removal of PFA, cells were washed with 1X PBS. Cell monolayers were then blocked for one hour in PBS with 3% BSA supplemented with 0.1% TritonX-100. After blocking, monolayers were incubated with CHIK-48 primary antibody that cross-react with Mayo virus E2 Envelope Glycoprotein (BEI resources; NR-44002 −1:500) overnight at 4 °C in blocking solution. Unbound primary antibodies were washed with PBS and incubated with Alexa-488-Conjugated secondary antibody (Invitrogen; A28175-1:500) for one hour at RT in PBS/3%BSA/0.1%TritonX-100. The secondary antibodies were removed and washed with PBS. Chips were imaged on an Olympus BX41 epifluorescent microscope, and foci were quantified.

### F) Biological Samples

The following reagent was provided by Centers for Disease Control and Prevention for distribution by BEI resources, NIAID, NIH: *Aedes aegypti*, Strain D2S3, Eggs, NR-45838. The following reagents were obtained through BEI Resources, NIAID, NIH: *Culex tarsalis*, Strain YOLO, Eggs, NR-43026, genomic RNA from Zika Virus, MR 766, NR-50085, and as part of the WRCEVA program: Mayaro Virus, BeAn343102, NR-49909. A hatching broth was prepared by grinding 1g of fish food granules (Aqueon goldfish granules) and mixing in 1L of DI water. The solution was autoclaved and allowed to cool down to room temperature. The broth was then added to a plastic tray filled with 1L DI water and eggs were hatched in the trays (200 eggs per tray). Ground fish food granules (Aqueon goldfish granules) we reprovided as feed for the larvae. The pupae were transferred to cups and placed in polypropylene rearing cages (Bugdorm, Taiwan). Mosquitoes were reared at 28 °C, 80% humidity with 12 hour light-dark cycles. Flasks with sugar water (10%) and cotton wicks were placed in the rearing cages for the mosquitoes. *Aedes aegypti* strain KPPTN, *Aedes albopictus* strain BP, and ZIKV strain FSS13025 were provided by Louis Lambrechts (Institut Pasteur). *Aedes aegypti* DNA was obtained using lab extraction from homogenized, dead mosquitoes.

### G) ZIKV infectious blood meal

Approximately 55 *Aedes aegypti* females (11 days old) housed in cardboard containers were offered an infectious blood meal of defibrinated sheep blood supplemented with Zika virus at 1 × 10^7^ FFU/mL using a hemotek blood feeder. Females were allowed to feed for 15 minutes and subsequently cold anesthetized to separate fed and non-fed individuals. Infected adults were maintained at 28C and 80% humidity having continuous access to a 10% sugar solution. Experiments were performed at 17 and 18 days post infection.

### H) Fluorescent saliva imaging

To facilitate imaging of fluorescent saliva, *Ae. aegypti* were allowed to feed on a solution containing 0.4% rhodamine B and 10% sucrose in water for at least 48 hours. The rhodamine B ingested during sugar feeding stains the mosquito body including saliva. [36] Imaging was performed using the biteOscope as described previously [21] with the following adjustments: A 532 nm, 5 mW laser (Sparkfun, COM-09906) was used for illumination, and a 550 nm longpass filer (Thorlabs FEL0550) was used as an emission filter. Fluorescent saliva droplets on vectorchips were quantified by scanning the chips before and after the biting assays on Typhoon FLA 9000 gel scanner.

## Acknowledgements

This project was supported by grants from United States Agency for International Development (USAID) and National Institute of Health (NIH). Part of this work was performed at the Stanford Nano Shared Facilities (SNSF), supported by the National Science Foundation under award ECCS-1542152, at Stanford University.

We thank Louis Lambrechts (Insect Virus Interactions Unit, Institut Pasteur, Paris, France) and Lark Coffey (School of Veterinary Medicine, University of California, Davis, CA, USA) for providing access to arbovirus laboratories. USAID, CDC, SNF, NIH, BEI.

FJHH was supported by a Rubicon fellowship for the Netherlands Foundation for Scientific Research, a Career Award at the Scientific Interface from the Burroughs Wellcome Fund, and a Marie Curie Fellowship from the European Union.

SP and JLR were supported by NIH Grants R01AI128201, R01AI150251, R01AI116636, USDA Hatch funds (Accession #1010032; Project #PEN04608), and a grant with the Pennsylvania Department of Health using Tobacco Settlement Funds.

