## Supporting Information File for "Vectorchip: Microfluidic platform for highly parallel bite by bite profiling of mosquito-borne pathogen transmission"

### Supporting Figures

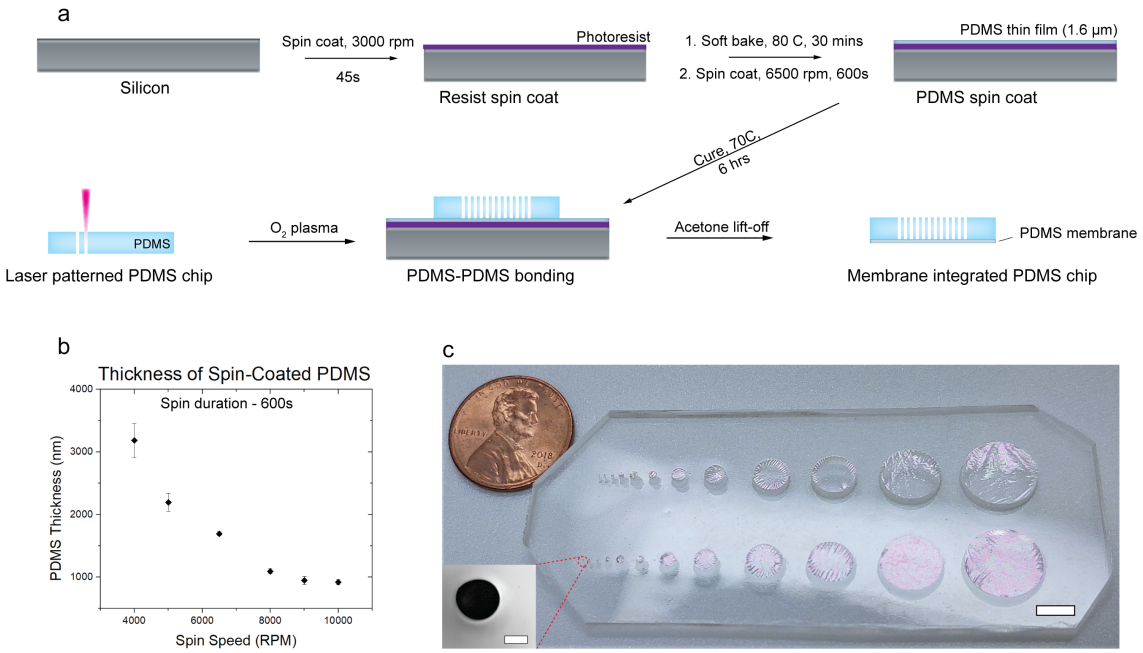


Figure 1: **Fabrication of vectorchips.** a) Schematic shows detailed steps for the fabrication of vectorchips. Laser patterning and soft lithography are used for fabrication. A sacrificial resist layer is spun onto a flat silicon wafer. A 30 min soft-bake was performed to dry the resist layer. A thin PDMS film is spin-coated onto the resist layer and placed in an oven for curing. PDMS blocks (Sylgard 184, 1:1 base:crosslinker) were prepared in lab and patterned using a $CO_2$ laser cutter. The patterned chips are bonded to the PDMS thin film using plasma oxidation. The wafers are placed in acetone where the resist layer is dissolved and the chips can be retrieved for cleaning and drying. b) Variation in thickness of the PDMS membrane with change in spin speed for a spin time of 600 seconds. c) A chip showing close to 2 orders of magnitude variation in well and membrane size. The smallest wells are 150 μm in diameter whereas the largest are 1 cm. Scale bars represent 5 mm in (c) and 100 μm in (c, inset).

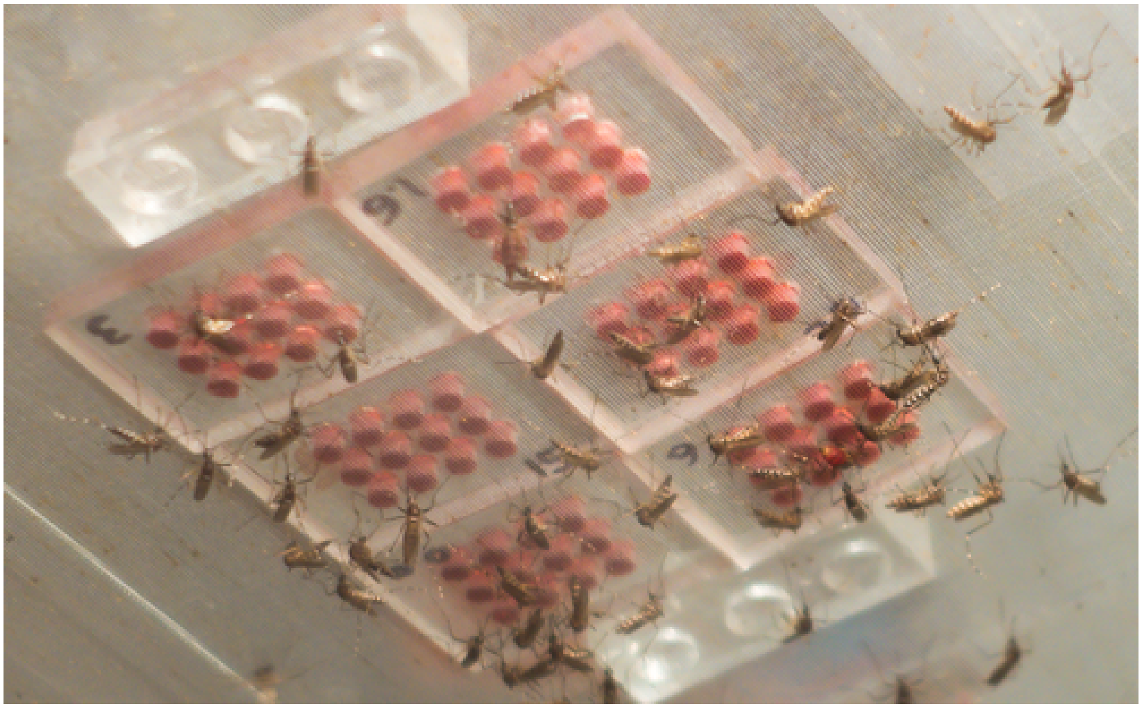

Figure 2: **Blood feeding on chips.** *Aedes aegypti* mosquitoes blood feeding from small chips with thin PDMS membranes.

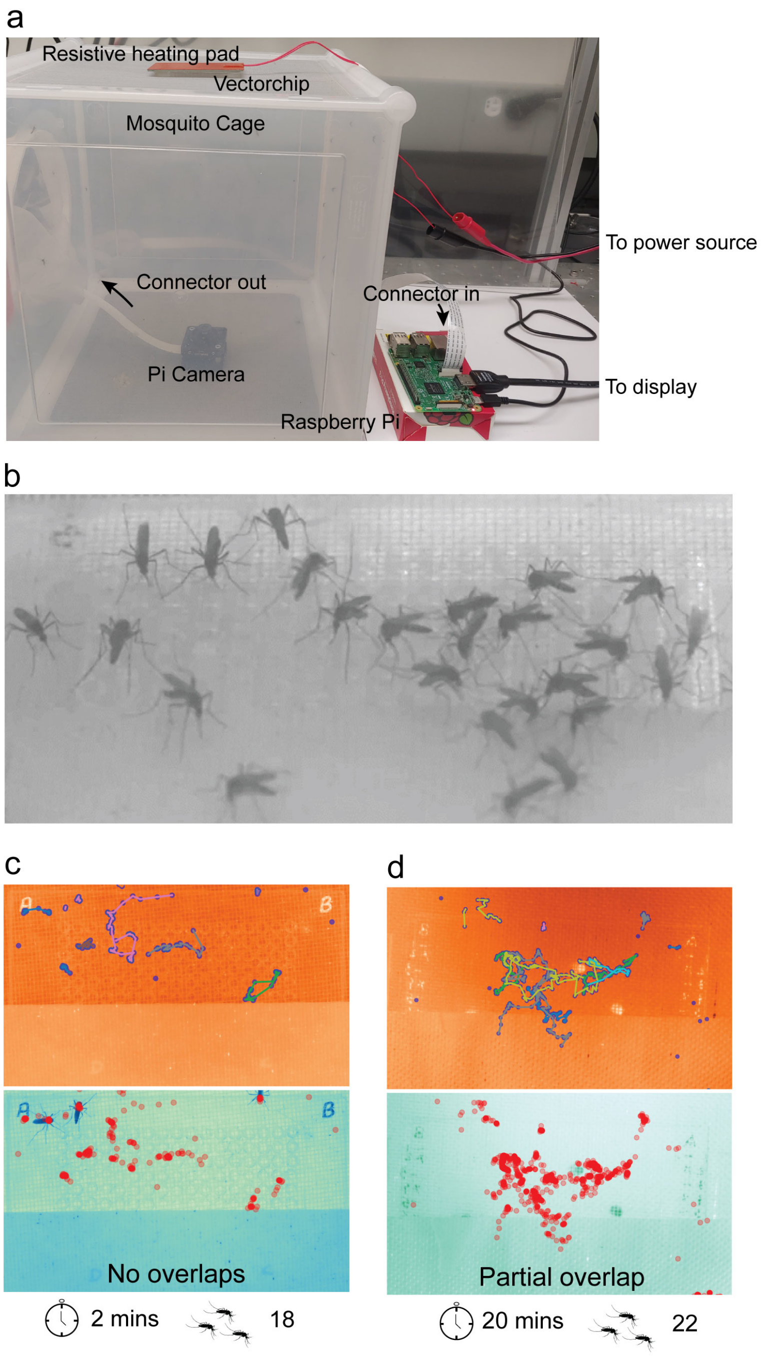


Figure 3: **Tracking on-chip biting.** a) Image showing setup for tracking mosquito movement during bite assays. A raspberry pi camera is placed at the bottom of the cage and mosquitoes are allowed to access the sugar water through the mesh barrier at the top of the cage. A resistive pad is used as the source of heat to attract the mosquitoes. b) Mosquitoes drinking sucrose from a vector chip. c, d) Tracking data showing two cases where no overlap or partial overlap between mosquito activity is observed dependent on the time of mosquito-chip interaction. Regions with no overlap in trajectories can provide statistical information about probing performed by individual mosquitoes.

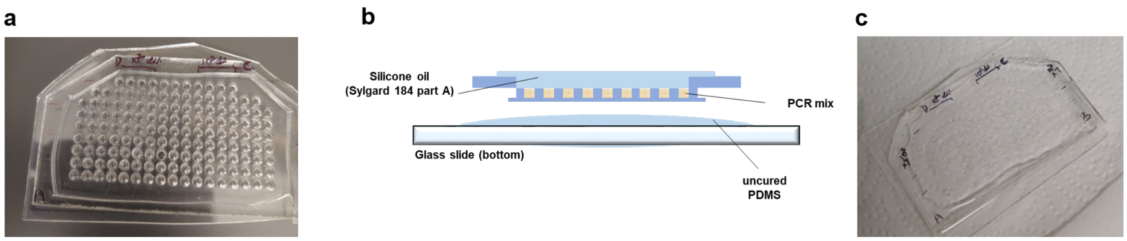


Figure 4: **PCR on-chip.** a) PCR reaction mix is pipetted into all the wells (volume - 4 µL per well) b) The top and bottom of the chip are covered with silicone oil to prevent evaporation. The bottom silicone is a 1:1 mix of Sylgard 184 base:crosslinker and therefore helps the chip adhere to the glassslide. The top silicone oil is Sylgard 184 base (without crosslinker). c) Chip with PCR reaction mix and silicone oil layers prior to PCR.

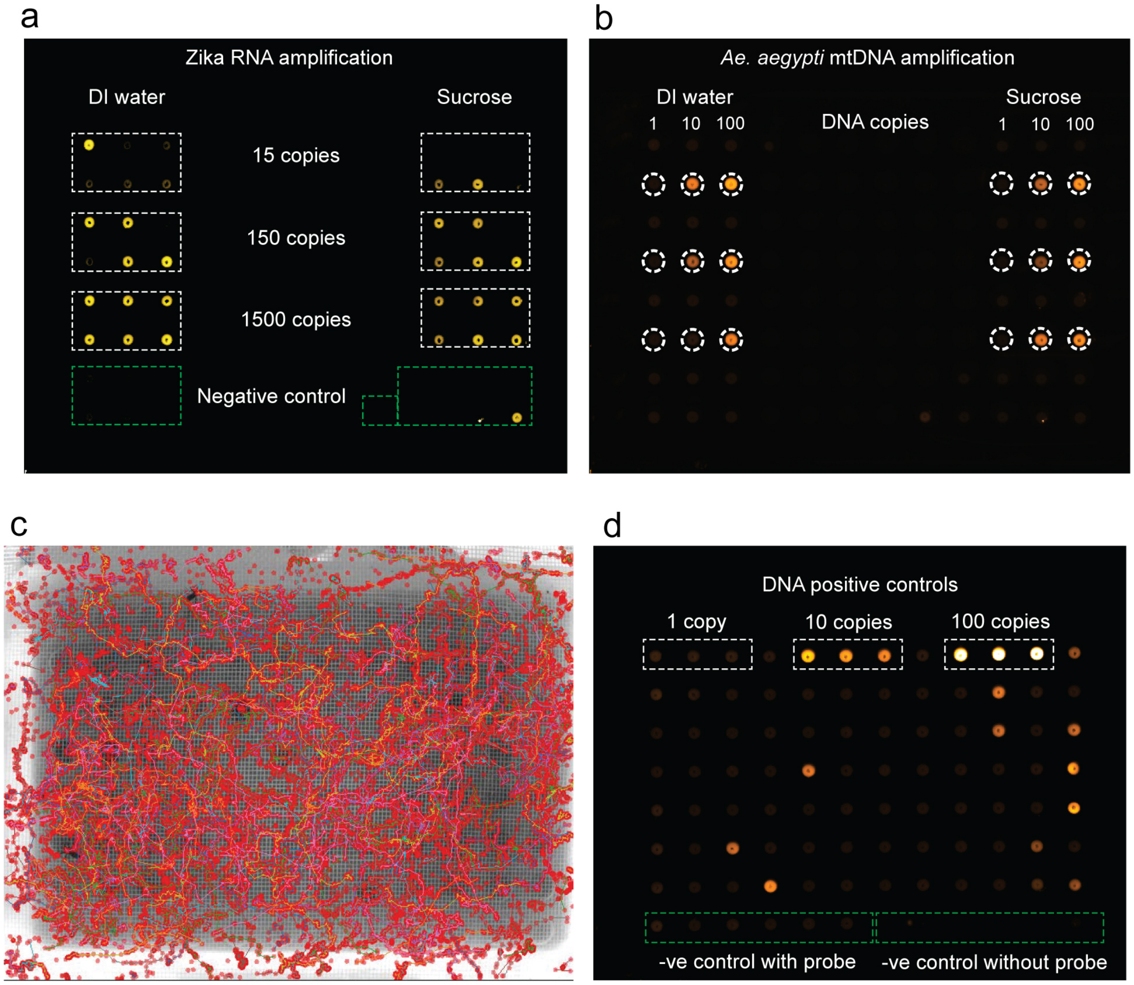


Figure 5: **PCR on 96 well plate.** PCR assays with manually spiked (a) RNA and (b) DNA was performed on a 96 well plate for 42 cycles (4 µL reaction volume). The reactions were performed in both DI water and 10% sucrose to test if sucrose inhibits the reactions. Amplification was observed in both cases for RNA (15-1500 copies) and DNA (10-100 copies). RNA amplification was less consistent for 15 RNA copies (5/6 wells showed increase in fluorescence in DI water, with weaker amplification indicated in 4 wells. 2/6 wells showed amplification in sucrose.) 1 false positive was observed for RNA amplification. (c, d) 96 well plate covered with parafilm is used to simulate the vectorchip experiments. Low melting point agarose at 2% concentration is used as feeding media and covered with a layer of parafilm. The parafilm is removed post bite as it is incompatible with thermocycling. The layer is removed carefully to avoid cross-well contamination. (c) On-chip tracking reveals mosquito translocation covering all regions of the 96 well plate. (d) PCR assays demonstrate detection of *Aedes aegypti* mitochondrial DNA in $\sim$ 17% of the available wells..

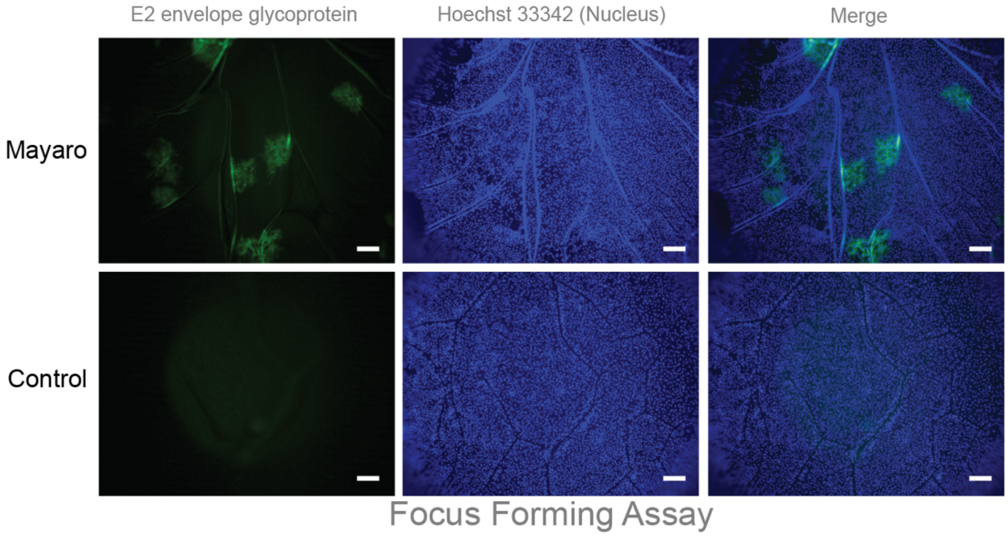


Figure 6: **Focus Forming Assays (FFA) on chip.** Mayaro viral particles were manually pipetted to sample wells in vectochips. A monolayer of vero cells are grown on the skin-mimic PDMS membrane (labeled using Hoechst 33342, nuclear stain). The Mayaro viral particles infect the cell monolayer resulting in foci and are visualized using a labeled antibody against the E2 envelope glycoprotein. The scale bar is 100 μms.
